# Shared resources between visual attention and visual working memory are allocated through rhythmic sampling

**DOI:** 10.1101/567602

**Authors:** Elio Balestrieri, Luca Ronconi, David Melcher

## Abstract

Attention and Visual Working Memory (VWM) are among the most theoretically detailed and empirically tested constructs in human cognition. Nevertheless, the nature of the interrelation between selective attention and VWM still presents a fundamental controversy: do they rely on the same cognitive resources or not? The present study aims at disentangling this issue by capitalizing on recent evidence showing that attention is a rhythmic phenomenon, oscillating over short time windows. Using a dual-task approach, we combined a classic VWM task with a detection task in which we densely sampled detection performance during the time between the memory and the test array. Our results show that an increment in VWM load was related to a worse detection of near threshold visual stimuli and, importantly, to the presence of an oscillatory pattern in detection performance at ∼5 Hz. Furthermore, our findings suggest that the frequency of this sampling rhythm changes according to the strategic allocation of attentional resources to either the VWM or the detection task. This pattern of results is consistent with a central sampling attentional rhythm which allocates shared attentional resources both to the flow of external visual stimulation and also to the internal maintenance of visual information.

## 1. Introduction

A key challenge for cognition is to efficiently process sensory information in order to guide behavior while also strategically allocating our resources based on recent experience and current goals. This interplay between externally (perceptual) and internally generated representations has been a core focus of research since the dawn of experimental psychology **(Perky, 1910)**. In this interplay, a central role has been attributed to both attention, i.e. the set of mechanisms that tune psychological and neural processing in order to identify and select the relevant events against all the competing distractors **(Nobre & Mesulam, 2014)**, and working memory, i.e the brain system maintaining and manipulating the information necessary for high level cognitive tasks **(Baddeley, 1992)**.

Both of these constructs have been extensively examined in the visual domain. This led on the one hand to the formulation of the concept of Visual Working Memory (VWM) as the active maintenance of visual information to serve the needs of an ongoing task **(Luck and Vogel, 2013)**. On the other hand, the concept of selective attention has encompassed all the ways the brain controls its own information processing **(Chun, Golomb & Turk-Browne, 2011)**. This broadness in the construct of attention has led some authors to advocate a distinction between external and internal attention, where the first refers to the selection and modulation of sensory information as it initially comes into the mind, in a modality specific representation, whereas the second includes the selection and modulation of internally generated information **(Chun et al. 2011)**. In this framework, VWM would be the nexus between the internal and external, as an interface through which attentional mechanisms select information from the external world which must be actively maintained as internal representations due to task relevance **(Chun, 2011; Chun et al. 2011)**.

Many theories of attention and working memory emphasize core differences between the two mechanisms. For example, in an extensive review of the literature Pashler **(1994)** suggests that the main cause of competition between attention and working memory tasks relies on a bottleneck limitation rising from impossibility to execute some types of processes simultaneously, rather than a shared resource. An even stronger claim comes from the work by Woodman, Vogel and Luck **(2001)** who suggest that the efficiency of visual search is not affected by the increase of load in a concurrent VWM task. However, in their study there was an overall and significant increase in reaction times (RT) for the visual search task in the presence of a VWM task compared to the condition without a concurrent memory task. More recently, Hollingworth and Maxcey-Richard **(2013)** showed that inserting a difficult search task during the maintenance of a set of items in VWM did not affect memory performance when attention was selectively deployed to a cued item. Nevertheless the overall performance was reduced in the dual task condition.

Another perspective suggests instead that both internal and external attention rely on shared supramodal attentional resources and on common neural substrates, which would result in a competition between the two **(Kiyonaga & Egner, 2013)**. In support for the idea of competition, there are several studies showing that discrepancies between content in VWM and visual targets impacts on performance in visual search **(Soto, Heinke, Humphreys, & Blanco, 2005)**, as well as in more basic phenomena related to attentional deployment such as saccade correction **(Hollingworth & Luck, 2009)** and saccade landing **(Hollingworth, Matsukura & Luck, 2013)**.

Moreover, the first part of this claim, i.e. the reliance of these two facets of attention on common neural substrates, is supported by converging neuroimaging evidence. For example, the increase in VWM load is related to activity suppression in the temporal parietal junction (TPJ) during the maintenance phase in VWM, and at the same time the increase in VWM load was related to lower detectability of task irrelevant stimuli **(Todd, Fougnie & Marois, 2005)**. Moreover, both the encoding phase of VWM and of a demanding visual search task share some common neural substrates including occipito-temporal cortex, intraparietal sulcus (IPS), precuneus, precentral sulcus, and the frontal gyrus **(Mayer, Bittner, Nikolić, Bledowski, Goebel, & Linden, 2007)**. In addition, IPS activity, traditionally considered part of the Dorsal Attentional Network involved in top-down, goal directed control of incoming information **(Corbetta & Shulman, 2002; Fox, Corbetta, Snyder, Vincent & Raichle, 2006)** was enhanced in a load-dependent fashion during the VWM maintenance phase, reaching a plateau at around 4 elements **(Todd & Marois, 2004)**. Multivariate Pattern Analysis (MVPA) has revealed as common substrate for crossmodal (verbal and visual) working memory storage in the IPS and bilateral frontal regions, where the neural activation patterns differentiating high-and low-verbal WM load can be predicted by neural patterns dissociating high and low load in a visual WM task, and vice versa.

Further support for a shared mechanism comes from EEG studies. An electrophysiological counterpart of the results obtained by Todd and Marois **(2004)** was reported by Vogel and Machizawa **(2004)**, who recorded Event Related Potentials (ERPs) from participants performing a VWM task and measured the so-called Contralateral Delayed Activity (CDA), a large negative component over the hemisphere contralateral to the set of items to be memorized. Interestingly, the amplitude of this component reached a plateau at approximately four items, it was correlated with individual VWM capacity and it has been subsequently linked to a very similar, negative component appearing contralateral to the hemifield in which the participants had to direct their focus of attention in a visual search task **(Emrich, Al-Aidroos, Pratt, & Ferber, 2009)**. In summary, the evidence outlined above suggests that some common top-down brain mechanism of control might subserve both the attentional modulation of perceptual information and the different phases of encoding, maintenance and retrieval in VWM **(Gazzaley & Nobre, 2012)**. A fundamental point that remains to be addressed, as also suggested by Kiyonaga and Egner **(2013)** is whether VWM competes with the attentional modulation of the incoming visual information. If so, at which level does this competition happen?

An influential point of view regarding the locus of competition is provided by the ‘Load Theory of Attention and Cognitive Control **(Lavie, 1995; Lavie, 2005; Lavie & Dalton, 2014)**. According to this theory, the degree of processing of visual distractors depends on the perceptual load of the task: with a low demand, an increased number of distractor would be processed, an effect reduced in high demanding perceptual tasks. In parallel, the theory advocates that target prioritization in the face of distractors would depend on the current availability of executive control functions for active maintenance of the processing priorities. When these functions are loaded by concurrent tasks, this would result in an increased processing of distractor stimuli (an opposite effect to the one postulated for the perceptual load). Several lines of evidence within the theoretical framework proposed by Lavie and colleagues indicate that in dual task conditions high VWM load would mainly burden the perceptual system. For example, when concurrently performing a delayed match to sample (DMS) and a detection task during the maintenance phase in VWM, high VWM load is accompanied by reduced detectability of near-threshold visual stimulus detection, and decreased activity in early visual cortices compared to low VWM load **(Konstantinou, Bahrami, Rees & Lavie, 2012)**. A similar effect on detection is obtained by taxing the perceptual system with a demanding visual search task **(Konstantinou, & Lavie, 2013)**, whereas a WM task oriented in taxing the executive system (digit span) would result in an increase in detectability of both irrelevant and relevant stimuli **(de Fockert & Bremner, 2011; Konstantinou, & Lavie, 2013)**.

The pattern of results described above would suggest that competition between attention to external incoming information and internally maintained visual information would take place at an early stage of the perceptual processing. However, what remains to be clarified is how this bottleneck is resolved. An analogous debate on division of attentional resources among external stimuli has historically seen two main positions: the first suggesting a “parallel” strategy implying that the limited resources would be spread across the items to be attended simultaneously **(McElree & Carrasco, 1999)**, and a second perspective supporting a “sampling” strategy, thus underlying a serial attentional shift focused for every single item to be attended **(Treisman & Gelade, 1980)**. A seminal study **(VanRullen, Carlson, & Cavanagh, 2007)** suggested that when attention is split among potential task-relevant locations, a sequential strategy of sampling is used and, moreover, this sampling of information might be a rhythmic phenomenon with a periodicity of ∼7 Hz **(VanRullen et al. 2007; Busch & VanRullen, 2010; VanRullen 2016)**. This idea has found further support in the work by Landau and Fries **(2012)** using an exogenous spatial attention cueing paradigm, which uncovered the existence of a fluctuation in detection accuracy at 4 Hz, counterphased for the two spatial positions. Such an oscillatory pattern suggests that our capacity to focus attention waxes and wanes periodically, and that such periodicity might be determined by the amount of competition among the items or the positions to be currently attended **(Landau & Fries, 2012; Fiebelkorn, Saalmann, & Kastner, 2013; VanRullen, 2018; Fiebelkorn, Pinsk & Kastner, 2018)**. If such oscillations occur when external attention is divided, for example, among several positions, and if attention involves a central mechanism that allocates a limited amount of resources among different channels of information both internal and external **(Kyonaga & Egner, 2013)**, then we can hypothesize the existence of an analogous oscillatory phenomena when the visual system must support both a demanding (external) visual detection task and a challenging (internal) VWM task.

The present study investigates this question by taking advantage of the methodology of temporal dense sampling of behavioural performance **(Fiebelkorn, Foxe, Butler, Mercier, Snyder & Molholm 2011; Benedetto & Morrone, 2017)**. We probed behavioural performance in the detection of a near threshold visual stimulus presented at many equally spaced stimulus onset asynchronies (SOA) while participants maintained in VWM a set of stimuli in order to perform a Delayed Match to Sample (DMS) task **(Luck & Vogel, 1997)**. We hypothesized that the absence of VWM load would result in an oscillation in detectability at approximately 7 Hz, as previously reported, whereas this fluctuation should shift toward slower frequencies (approximately 4 Hz) with the gradual increase of VWM load in line with the idea of division of attention between competing representations. We expected also to reproduce the effects shown by Kostantinou and collaborators **(2012, 2013)**, according to which a decrease in the detectability of a near threshold stimulus would be expected when the perceptual system was taxed with a VWM task, at different load conditions, compared to the absence of VWM load. Finally, given the great amount of literature reporting individual differences in VWM capacity, linked to differences in electrophysiological response **(Vogel & Machizawa, 2004; Emrich et al. 2009)**, detection capacity in a dual task setting **(Kostantinou et al. 2012; Kostantinou et al. 2013)** and in the cortical thickness and surface of primary visual cortex cortices **(Bergmann, Genç, Kohler, Singer & Pearson, 2016)**, we hypothesized that individual differences in the strategic deployment of attention during the dual task would also be reflected in the rate of temporal fluctuations in performance, leading to different oscillatory patterns in the detection task between low-capacity and high performing participants in the VWM task.

## 2. Methods

### 2.1 Participants

Twenty-two participants took part in the experiment (12 females, mean age: 24.4, SD=3.5 years). All participants reported normal or corrected-to-normal vision and normal hearing and gave informed written consent. The experimental protocol was approved by the University of Trento ethical committee and was conducted in accordance with the Declaration of Helsinki.

### 2.2 Apparatus and stimuli

The experiment was programmed using PsychToolbox 3.0.14 running under MATLAB 2015b, and ran on a HP compaq 8000 Elite CMT computer. Stimuli were displayed on a VIEWPixx/EEG 22” screen with refresh rate of 100 Hz. The luminance of the room was measured with Konica Minolta illuminance meter T-10, yielding a value of 0.8 lumex in the room and 7 lumex at the front of the screen.

The target stimulus (hereafter referred to as “flash”) in the detection task consisted of a circular, luminance defined Gaussian blob with a diameter of 0.5° presented on a grey background (RGB: 128, 128, 128). The luminance of the blob with respect to the screen was adjusted according to a QUEST staircase procedure performed before each experimental session, and resulted in an average increment of luminance of 7.48% (SD=1.2%), computed across participants and sessions. The subjects were instructed that the flash might appear in variable positions, but always in an area delimited by a circle in the center of the screen (see Figure 1) with a diameter of 4°. The flash always appeared within a radius of 1.5° from center, in order to prevent overlapping or facilitatory effect due to enhanced contrast between the flash and the circle on screen (RGB: 0,0,0). The items to be memorized consisted of 4 squares, length of 4°, distant 2.8° from the center of the screen, each one presented in a separate quadrant of the screen. They were assigned one of the following RGB triplets: red (198,0,0), purple (191,0,198), blue (0,0,198), light blue (0,191,198), green (33, 198, 0), yellow (191, 198,0). The disappearance of the items from the screen was accompanied by an auditory stimuli used as “reset” (see details below), which was a sinusoidal 500 Hz sound presented for 20 ms through professional headphones. The rationale for the “reset” stimulus is that in dense sampling paradigms some event is necessary to align the timepoints of stimulus presentation to an ideal time zero (t_0_) point (**Fiebelkorn et al. 2011, Landau & Fries, 2012**). This reset event at t_0_ should be capable of temporally aligning the phase of rhythmic brain activity across trials in order to achieve a sufficient level of intertrial phase coherence, which is a fundamental step to characterize the underlying behavioural rhythms. Several previous studies have shown that both auditory and audiovisual stimuli are capable to generate such a phase reset (**Lakatos, Karmos, Mehta, Ulbert, & Schroeder, 2008; Fiebelkorn et al. 2011; Ronconi & Melcher, 2017; Ronconi, Busch & Melcher, 2018**).

**Figure 1:**
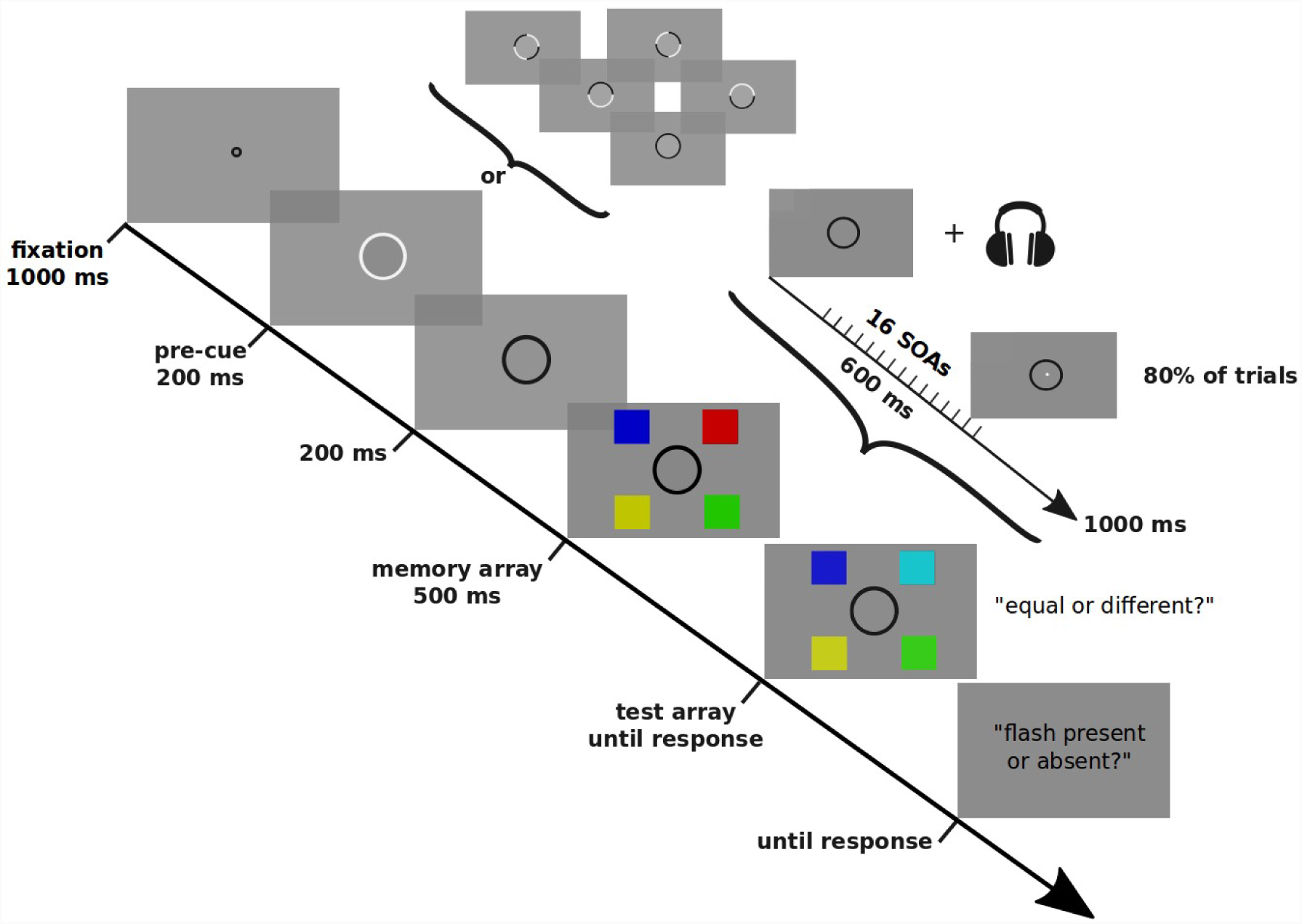
Schematic representation of the task procedure. Every trial started with a fixation period (1000 ms) followed by a pre-cue (200 ms). This could be either a white circle (main figure) a combination of different white quarter of circle (small screens north-east to the main pre-cue display all possible combinations) indicating the positions to be attended. The pre-cue was followed first by a black circle, on screen for the rest of the trial, and then by four coloured squares (memory array, 500 ms). The subject was instructed to remember selectively the items in the pre-cued positions. The disappearance of the square was accompanied by an auditory reset stimulus (see Methods). After 150 ms, in 80% of the trials, the flash stimulus appeared in the area delimited by the circle on screen, at 16 different, equally-spaced SOAs, up to 750 ms from the reset stimulus. After 250 ms a test array, displaying 4 coloured squares in which one of the pre-cued items could change in colour in 50% of the trials appeared, probing the participant to produce a judgment of equality (Equal vs Different) to the previous memory array. Once this first response was given, the participant was probed to report detection of the flash (Seen vs Unseen).

### 2.3 Experimental procedure

Every participant took part in two experimental sessions on two different days. At the beginning of each session participants were tested with a staircase procedure (50 trials) for determining their absolute contrast threshold in detecting the flash stimuli which were used in the main experiment. Afterwards, each participant completed the first session, composed of a practice block (20 trials) and by the real experiment (720 trials), which was divided in 5 smaller blocks of 144 trials each to prevent fatigue. The second session was identical, except that there was no practice block. Hence, each participant performed 1440 trials overall across the two sessions. Due to possible lack of familiarity with the task, the practice blocks were repeated when necessary in order to achieve a hit rate between 50% and 80% for the flash detection task. Both the practice trials and the real experiment had the same features, with the only difference that in the former a written colored feedback (“Correct!” in green, “Wrong…”, in red) on performance was provided at the end of each response while no feedback was given in the main experiment.

An outline of the task is provided in Figure 1. Each trial began with a fixation point of 1000 ms, followed by a pre-cue indicating the spatial positions to attend in the subsequent VWM task. This cue, circular in shape, could either indicate 0 positions (simple black circle), 2 positions (black circle with two white quarters of arc) or 4 positions (complete white circle, as in the figure) and lasted for 200 ms. It was necessary to use the precue paradigm in order to vary VWM load while also keeping visual stimulation constant across the different dual task conditions and avoid the possibility that the difference in load (0 to 4) was confounded with a difference in the amount of visual stimulation (and, potentially, the degree to which the display might reset and align any fluctuations in performance). Through this manipulation we were able to present 3 different VWM load conditions, from hereafter: load0, load2 and load4. In order to prevent possible confounds due to lateralized distractor suppression in VWM (**Sauseng et al, 2009; Wolff, Jochim, Akyürek & Stokes, 2017**), in the load2 the cued positions always included one quadrant on the right and one quadrant on the left hemifield. The cue was followed by a simple black circle without any other stimuli for 200 ms: notably the circle itself remained on screen for the whole duration of the trial, until the probe for response, since it delimited the area of possible appearance of the flash to be detected. After this blank phase, 4 coloured squares (in any load condition) appeared for 500 ms, and the subject was asked to memorize only the items in the cued position. The disappearance of the squares happened simultaneously with the auditory reset stimulus, after which, at 16 different equally spaced values of SOA (150 ms to 750 ms from the memory items disappearance and the auditory reset stimulus, in steps of 40 ms) the flash appeared in 80% of trials. After 1 second from the auditory reset stimulus, a new set of coloured squares appeared on screen in the same positions as before, with the only exception that one of the squares in one of the cued locations could change its color on 50% of trials; this set of items was accompanied with a probe to response (equal or different), and stayed on screen until response. Lastly, another screen probed the participant on whether the flash was present or absent.

### 2.4 Data Analysis

Data analysis was performed using custom scripts in MATLAB 2016a. Behavioural performance was first collapsed for each participant across the two sessions. Then we computed the accuracy in the VWM task and then trials in which the memory task was not performed correctly (on average, 8.45% of all trials) were excluded from further analysis for flash detection performance (**Kostantinou et al., 2012**). Subsequently, Hit Rate (HR) was computed for each temporal bin, for each memory load condition (**Macmillan & Creelman, 2004**). This resulted in 3 HR time series (one for each memory load) for each participant. In line with previous studies, these time series were detrended using a second-order polynomial (**Fiebelkorn et al., 2013**), and then Hanning tapered and zero-padded (**Landau & Fries, 2012**) in order to allow for better sensitivity to regular fluctuations in the data. After this pre-processing of the raw data, as a first analysis approach a Fast Fourier Transform was performed on the HR time course for each participant in order to obtain the amplitude for each frequency. The grand average of the spectra across participants was performed to obtain a single averaged spectrum for each load condition.

As a second analysis approach, we used a fitting procedure to determine the goodness of fit of the time series both at the individual and at the group level. Specifically, the adjusted R^2^ was computed from the fitting of a sinusoid using the following formula, where amplitude (A), frequency (*ω*) and phase angle (*θ*) were set as free parameters:

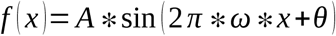

In order to test significance both in spectral analysis and in fitting analysis at the population level we performed nonparametric permutation tests by randomizing the hits and misses across different SOAs, separately for each participant and each condition. We conducted the same preprocessing steps described above also on the randomized data. After 10000 iterations for the spectra computation and after 1000 iterations for the fitting procedure, we constructed randomly sampled distributions either of amplitude values (in the case of spectra) or of adjusted R^2^ (in the case of fitting on grand averages. To test significant periodicities in the spectral analysis we individuated the peaks in the spectra exceeding the 95° percentile of the permutation distribution. In order to control for multiple comparison, the same test was repeated after selecting the peak of the 95° percentiles of the surrogate datasets showing the highest value over a range of frequencies of interest (**Nichols & Holmes, 2002; Huang, Chen, & Luo, 2015; Re, Inbar, Richter & Landau, 2019**), individuated between 2 and 10 Hz by our hypotheses (which would target phenomena in the classic theta range, between 4 and 7 Hz).

To avoid discarding possibly useful information embedded in the trend removed from the time series, we performed an analysis of the coefficients of the fitted second order polynomials defined by the following formula:

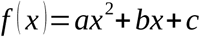

where the sign of the second order derivative of the function indicates whether the function has an upward (*f’’* (*x*)>0) or downward (*f’’* (*x*)<0) concavity. So, the second derivative of the function will result in:

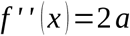

Which means that the sign of *a* describes the concavity of the parabola. In addition, its absolute values describes the steepness of the parabola itself. Both of these aspects were used to characterize the parabolic trends within the data.

HR and False Alarm rate (FA) were also used, collapsing across time bins, to compute the observers’ sensitivity (*d’*) and response bias i.e. criterion, for the 3 load conditions as follows (**Macmillan & Creelman, 2004**):

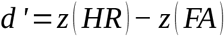

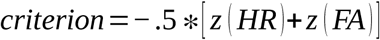

where *z* (*p*) represents the z score for a proportion *p* defined between 0 and 1.

Concerning the VWM task, a widely used estimate for the number of items currently maintained in memory in a delayed match to sample task is the so-called Cowan’s *k* (**Cowan, Elliott, Saults, Morey, Mattox, Hismjatullina & Conway, 2005**), defined as:

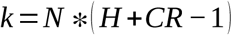

Where k is an estimate of the capacity in units of items, N is the number of items to-be-remembered, H is Hit Rate (measured as the probability to detect a change when the change is present) and CR is Correct Rejection, hence the probability to judge as different two arrays that are actually different.

## 3. Results

### 3.1 Signal Detection Theory (SDT)

As first step we computed typical measures of the SDT framework (see Methods) for detection of the near threshold stimulus in each VWM load condition (Figure 2). As expected, detection performance was better, with both higher HR and lower FA rates, in the zero VWM load condition **(Kostantinou et al. 2012; Kostantinou et al. 2013)**. A repeated-measures analysis of variance (ANOVA) with the within-subjects factor ‘load condition’ (load0, load2, load4) was first conducted on the Hit Rate (HR), showing a significant effect (F_(2,21)_ =10.09, p<.001, *η*^*2*^=.07). Post-hoc comparisons show differences between load0 and load2 (t_(21)_= 4.80, p<.001, *d=*.53) and between load0 and load4 (t_(21)_=3.34, p=0.003, *d*=.58), whereas no significant difference was detected between load2 and load4 (t_(21)_=1.05, p=.31).

**Figure 2:**
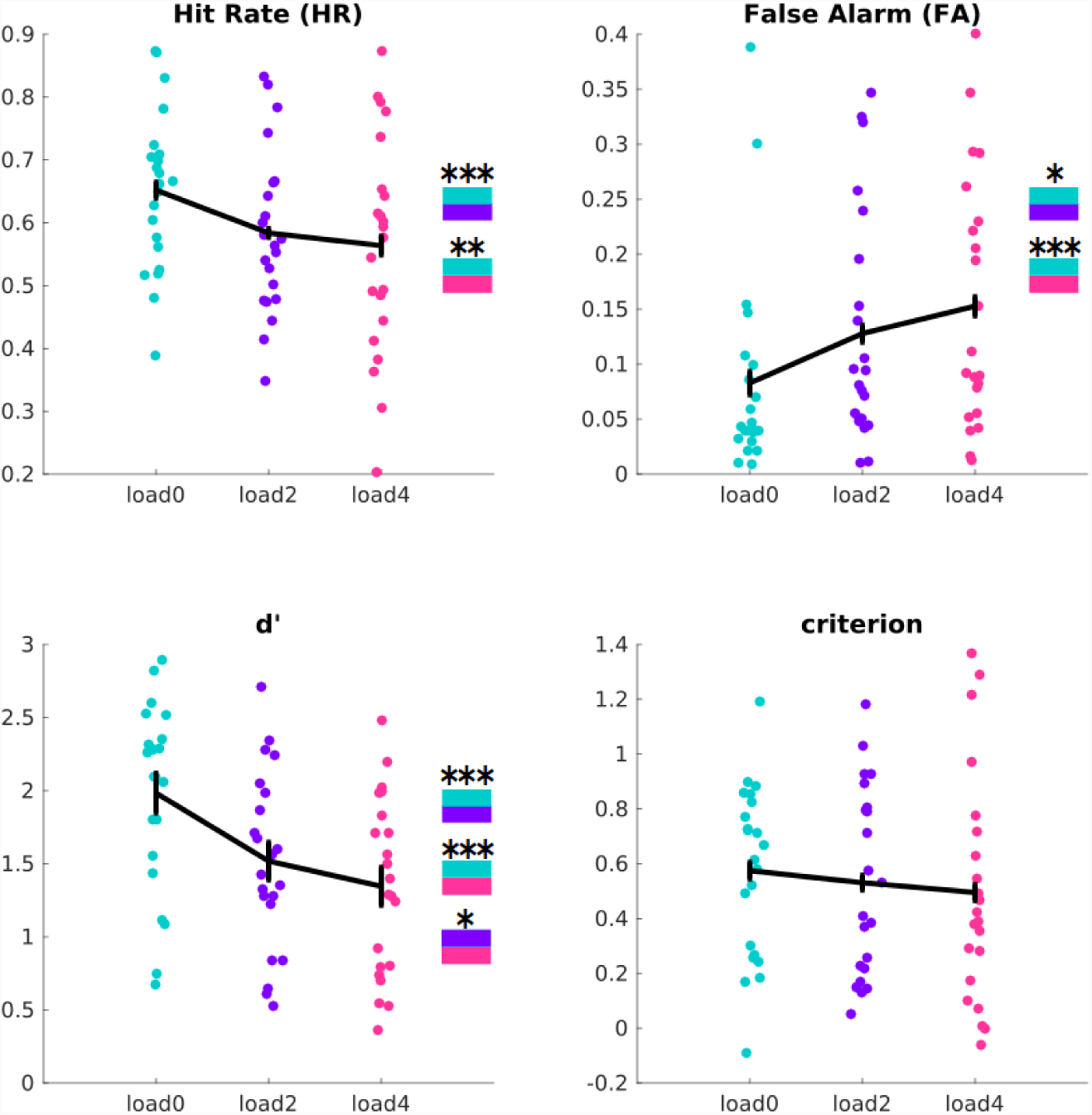
Signal Detection Theory results for the flash detection task: single dots show single participants in each load condition, whereas the errorbar displays within subject Standard Error of the Mean (SEM). Post-hoc comparison significant results are shown in a color coded fashion (turquoise load0, violet load2, magenta load4) in the boxes near the graphs; significance codes: * p<.05 ** p<.01 *** p<.001.

The ANOVA on False Alarm (FA) rate also revealed a significant effect of load condition (F_(2,21)_ =10.10, p<.001, *η*^*2*^*=.*07), and post-hoc comparisons showed that there were significant differences both between load0 and load2 (t_(21)_=-2.72, p=.012, *d*=-.44), and load0 and load4 (t_(21)_=-3.96, p<.001, *d*=-.66). There was a non-significant trend toward higher FA rate in the load4 compared to load2 condition (t_(21)_=-1.95, p=.063, *d*=-.22).

A similar ANOVA on *d’* showed a significant effect of load (F_(2,21)_ =22.66, p<.001, *η* ^*2*^ *=*.16), with post-hoc comparisons indicating significant differences between load0 and load2 (t_(21)_=5.41, p<.001, *d=*.75), between load0 and load4 (t_(21)_=5.19, p<.001, *d=*1.02) and between load2 and load4 (t_(21)_=2.16, p=.0418, *d=*.28). The ANOVA on the criterion did not show a significant effect of load condition (F_(2,21)_=1.27, p=.29). Overall, these results indicate that detection sensitivity was lower for the load conditions (load 2 and load 4) compared to the no load condition (load0), a result well explained by differences in HR and FA.

### 3.2 SDT and “good” vs “poor” performers

To target differences in parameters of SDT between individuals with high and low VWM capacity, we first computed Cowan’s K (see Methods) in both load conditions: load2 condition resulted in an average K_(2)_=1.72 (SD=.24), whereas load4 condition resulted in an average K_(4)_=2.52 (SD=.69).

We first divided the participants into two subsamples based on a median split with respect to the median of K_(4)_ (2.59) in the load4 condition. We refer to the median split in terms of “poor” or “good” performers with VWM in the dual task situation, where the first subsample (“poor”) was composed of those participants showing K_(4)_<2.59, and the second subsample of the remaining participants (“good”, N=11), respectively. The nomenclature focused on performance rather than capacity has been chosen since the VWM task was always in the context of a dual task. This means that we cannot infer whether this reflects a difference in VWM capacity in the absence of the second task. After this differentiation, we computed HR, FA, *d’*, and criterion for both groups (Figure 3).

**Figure 3:**
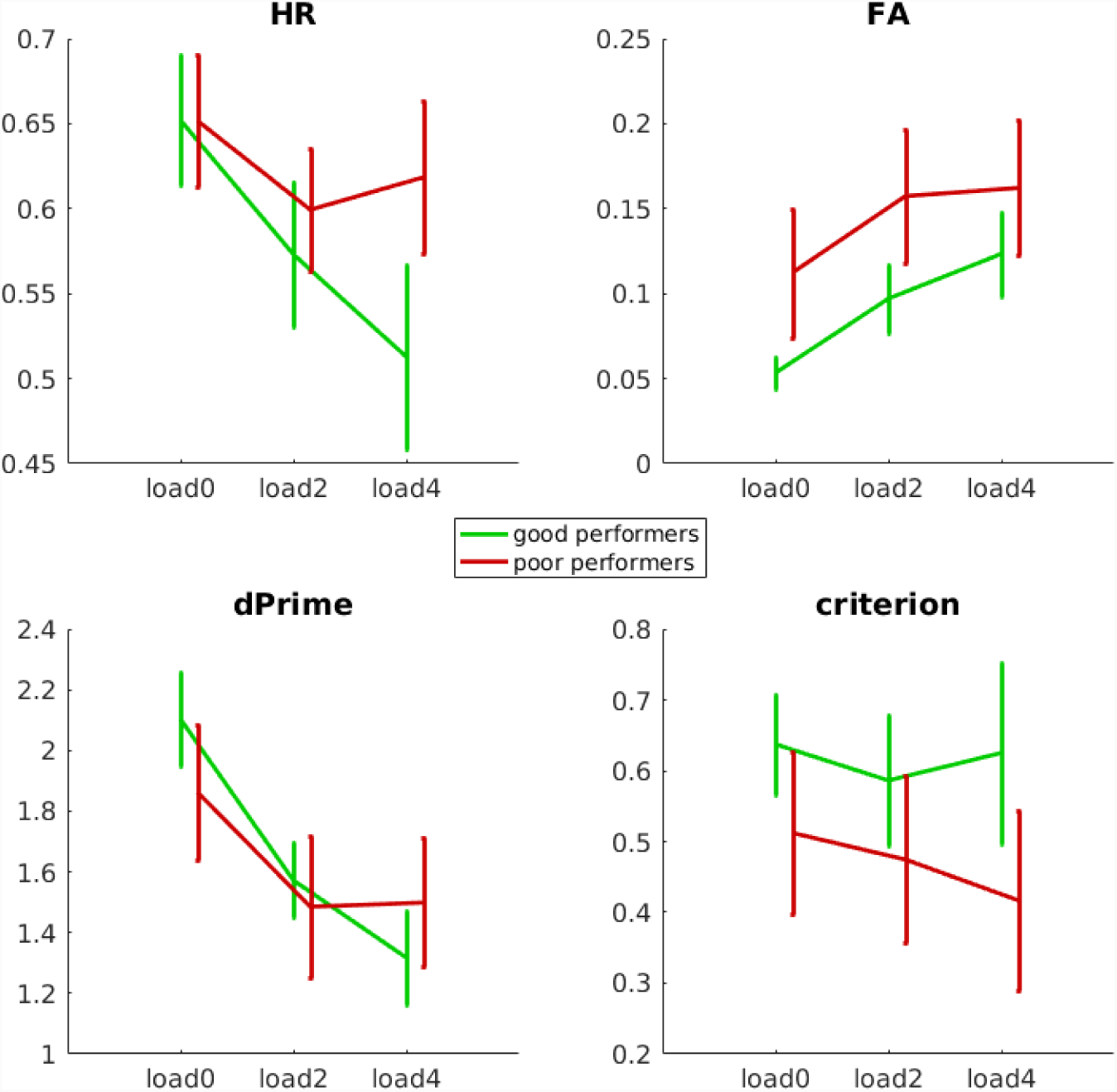
SDT results for the flash detection task, splitted between good and poor performers in the VWM task: green lines show “good” performers, red lines show “poor” performers, error bars display SEM.

For each parameter, similarly as before, we performed 2-way mixed model ANOVAs with the within-subjects factor memory load (load0 vs load2 vs load 4) and with the between-subjects factor group of performers (“good” vs “poor”). The ANOVA on HR confirmed the effects of load (F_(2,40)_ = 11.59, p<.001, *η*^*2*^*=.*07) and added a significant interaction between load and group (F_(2,40)_ =4.11, p=.023, *η*^*2*^*=.*02) but no significant effect of group emerged (F_(1,20)_= 0.515, p = .481). This interaction shows that “good” performers in VWM showed a drop in HR in load 4 compared to “poor” performers, whose performance in flash detection remained approximately stable between load 2 and load 4. This pattern of results was slightly different for the ANOVA performed on FA, confirming a significant effect of load (F_(2,40)_=9.69, p < .001, *η* ^*2*^ *=*.07) but no significant effect of group (F_(1,20)_ =1.89, p = .184) or interaction between group and load (F_(2,40)_=0.15, p = .861). A similar ANOVA on the d’ confirmed the significant effect of the load condition (F_(2,40)_ =23.68, p < 0.001, *η*^*2*^*=*.17). No significant effects in group condition (F_(1,20)_=0.1, p=.757) and interaction (F_(2,40)_=1.93, p = .158) were obtained. Also for the ANOVA on the criterion the main effects of load (F_(2,40)_=1.26, p=.294) and group (F_(1,20)_=1.19, p=.288), as well as their interaction (F_(2,40)_=.83, p = .443) were not significant.

### 3.3 Second order trend evaluation according to load conditions and group of performers

Before analyzing the fluctuation of HR in detection over time, we performed an analysis of the coefficient of the second degree polynomial (see Methods) fitted on each HR time series for each participant in each load condition. This analysis aimed at a better characterization of the evolution of HR over time. Figure 4 shows the single trends and the average trends for the two subgroups of “good” and “poor” performers in the VWM task.

**Figure 4:**
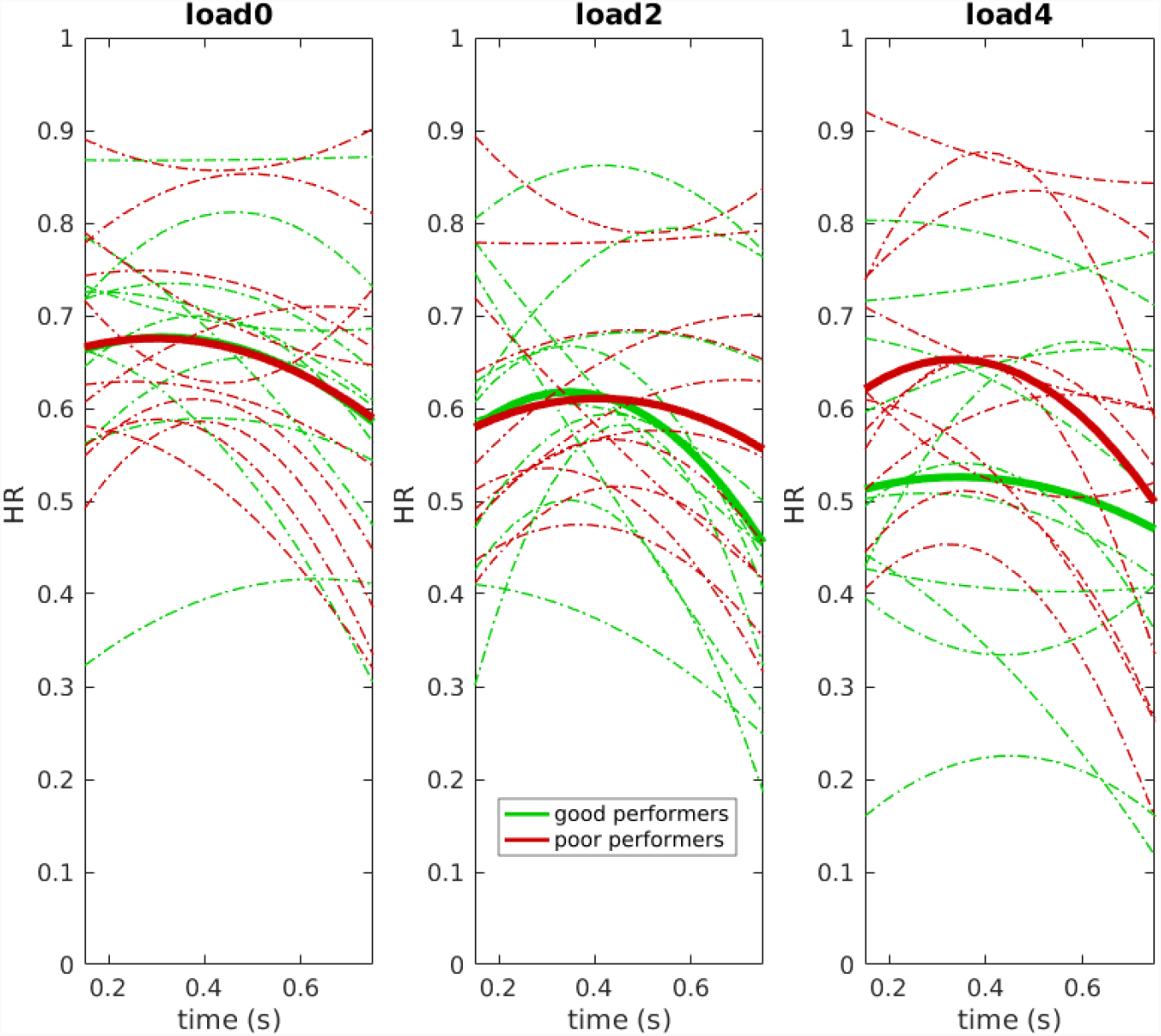
Second order (parabolic) trends between good and poor performers in the VWM task: green lines show “good” performers, red lines show “poor” performers, dashed lines show single participants and thick continuous lines show the average. It is possible to notice that most of the single subject parabolas and all the averaged trends show downward concavity (see main text). Furthermore the shape of the average parabola changes in the interaction between group and VWM load.

To assess concavity of the parabolas in each load condition we performed a non parametric, two-tailed sign test on *“a”* coefficients of the fitted 2nd order polynomials: in all the three load conditions the sign test was significant (load0, p=.008; load2, p=.008; load4, p=.026), suggesting that the concavity was consistently downward across the sample. To further elucidate the role of *“a*” coefficient in delineating the trend according to load (0, 2, 4) and group of performers (poor vs good), we performed a 2 way mixed model ANOVA with load as within-subjects factor and group as between-subjects factor. Even if no significant effects of group (F_(1,20)_=.002, p=.966) or load (F_(2,40)_=1.01, p=.373) were observed, a significant interaction between load and group emerged (F_(2,40)_ =3.86, p=.029, *η*^*2*^ *=*.09). This interaction can be fully appreciated in the average trends for each group as a function of each load condition, as shown in Figure 4. In particular, regardless of load condition, flash detection capacity over time was reduced when it was presented close in time to the memory array and test array (i.e. at the beginning and at the end of the dense sampling period, the two tails of the parabola), as displayed by the downward concavity. Furthermore this trend was influenced by the interaction between load and performer group, suggesting a steeper decay for poor performers while getting close to the memory array in the highest load condition, an effect present for good performers in load2.

### 3.4 Spectral analysis and sinusoidal fitting at population and at single subject level

As described above, one of the main aims of the study was to look for an oscillation in performance in the detection task, using the dense sampling approach. Figure 5 shows the average HR time series (before and after detrending), and the average spectra of individual participants in each load condition. To assess significance in spectra peaks we ran a non-parametric permutation test, and then corrected for multiple comparison the threshold for significance (see Methods). As can be seen in Figure 5 the only condition showing significant periodicities was load4, at ∼5 Hz (p=.044). The other two conditions show peaks similarly at ∼5 Hz for load0 and ∼4.16 Hz for load2, but without reaching significance (p=.190 and p=.423, respectively).

**Figure 5:**
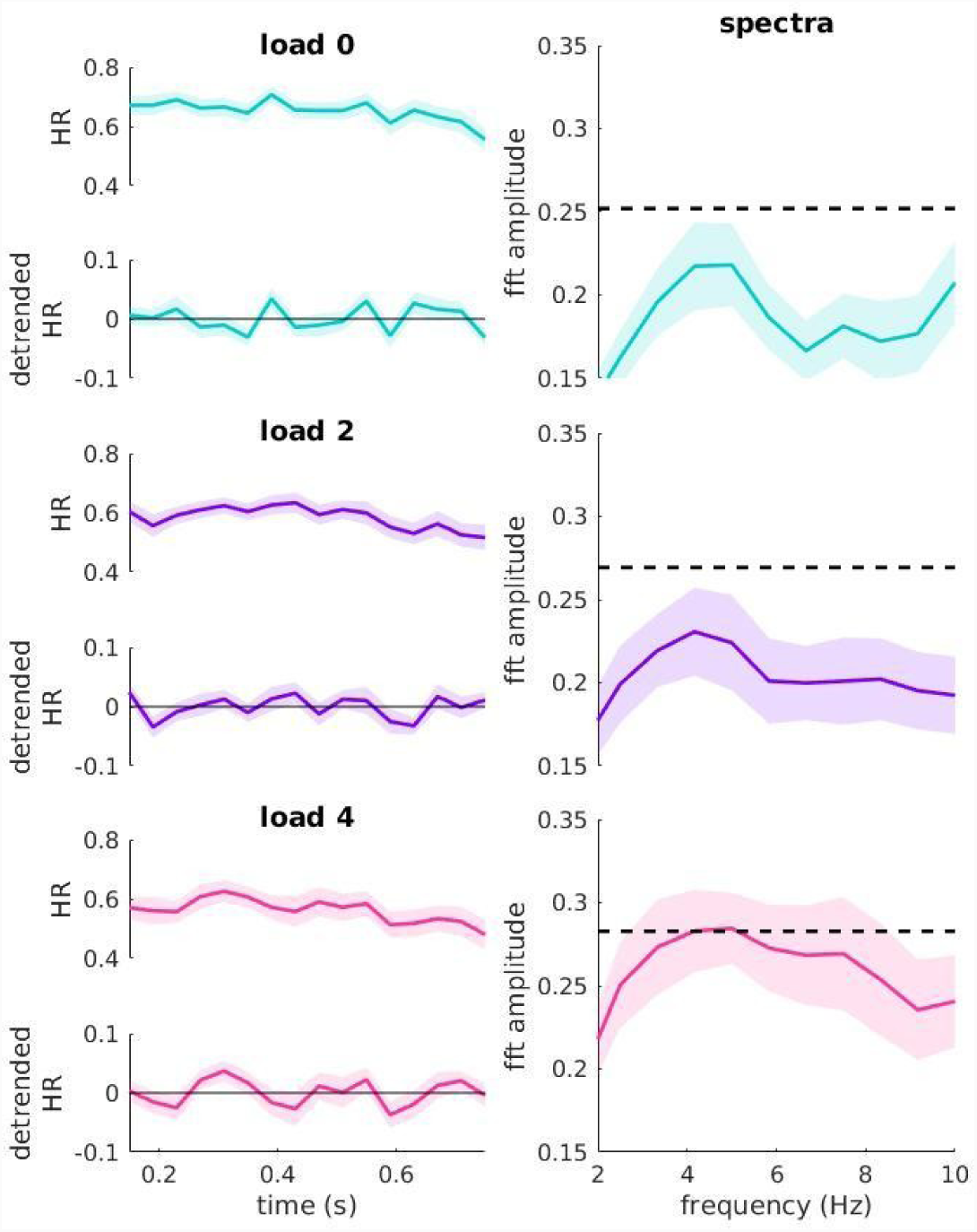
HR time series before and after detrend, and corresponding average spectra: shaded areas show SEM, colors are coded as in Figure 2, (turquoise load0, violet load2, magenta load4), black dashed line in the spectra insets show the permutation based threshold for significance corrected for multiple comparisons (see Methods).

To seek further support for the results obtained with the spectral analysis, we performed sinusoidal fitting (see Methods) both at the population level (on the grand average of detrended HR time series) and at the single subject level. At the population level (Figure 6) significance was again tested by the use of nonparametric permutation tests (see Methods), and the periodicity in load 4 was confirmed (best frequency fitted: ∼5.08 Hz, *adj R*^*2*^: .69, p=.014). Small discrepancies in the frequency fit with respect to the spectral analysis might be explained by the reduced frequency resolution of the spectral analysis itself. For load0 and load2 the best frequencies fitted on the grand average of HR detrended time series showed discrepancies with respect to the peaks in the spectral analysis: load0 showed the best fit at ∼7.08 Hz (*adj R*^*2*^ = .14) and load2 at ∼7.58 Hz (*adj R*^*2*^ = .3), even though neither load0 or load2 reached significance (p=.742 and p=.391, respectively), as shown in Figure 6.

**Figure 6:**
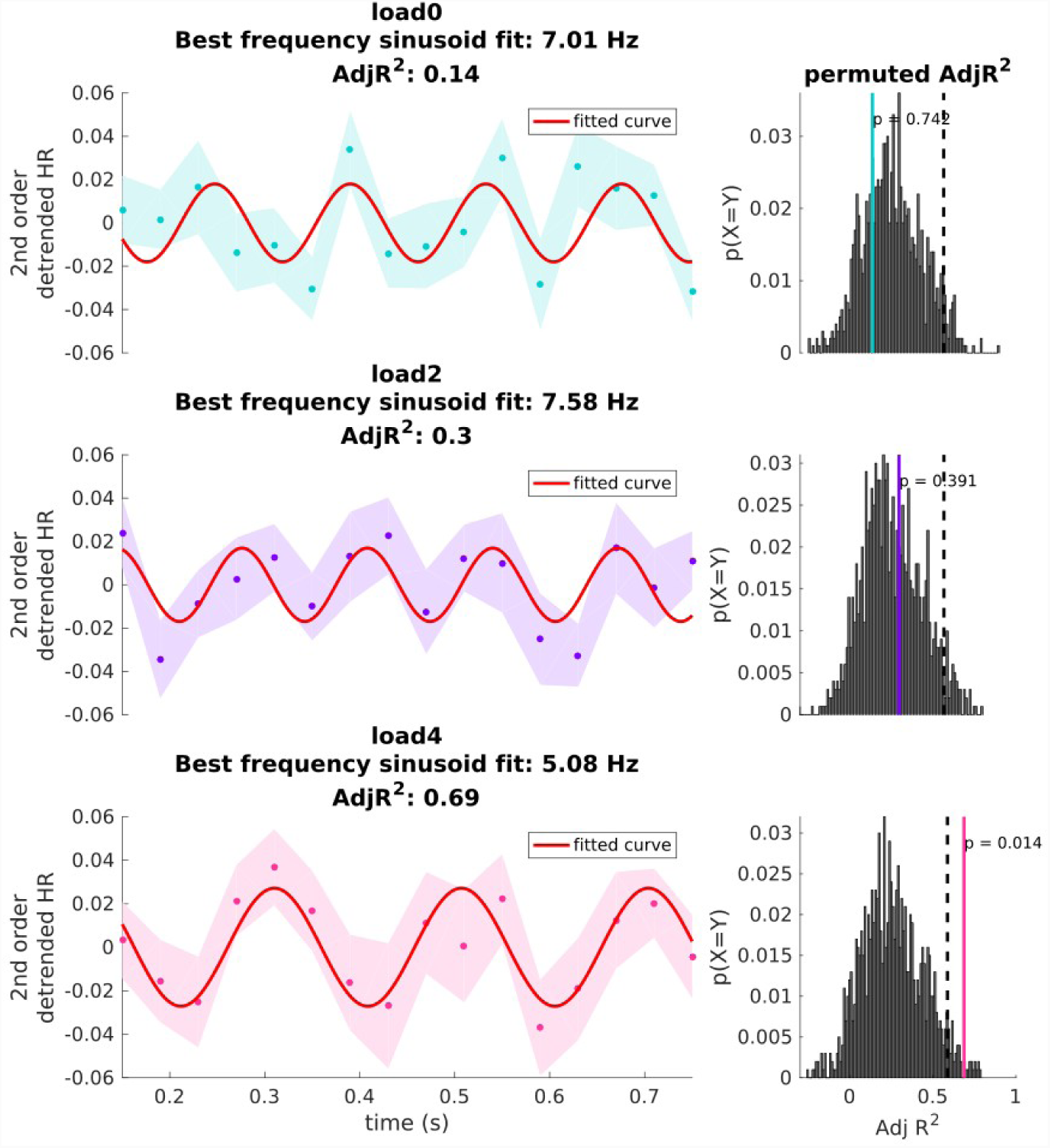
Average of HR time series after detrend with related sinusoidal fitting and significance testing: left insets show average trends, shaded areas show SEM, colors are coded as previously, (turquoise load0, violet load2, magenta load4), and the red line show the sinusoidal function fitted on data. The right insets show histograms of *adj R*^*2*^ obtained by the permutations in each specific condition, the black dashed line show the 95% percentile of the permuted distribution as threshold for significance, whereas the colored lines show where the observed *adj R*^*2*^ stands within the distribution of permuted values.

We performed an analogous fitting procedure to characterize the best fitting frequencies at the single participant level. Results are shown in Figure 7 and it is possible to note that the mean of fitted frequencies for load4 is at 5.88 Hz (median= 5.06, SEM = .55) whereas in the other two conditions a better average fit at higher frequencies was observed (load0 mean= 7.94, median = 8.00, SEM = .61; load4 mean = 7.87, median = 8.32, SEM = .71). Repeated measures ANOVA showed that this difference in best fitting frequencies across conditions was significant (F_(2,21)_= 3.57, p=.036, *η* ^*2*^ *=*.10), suggesting that when analyzing HR time series at the level of single participants the highest VWM load condition (load4) was associated with a slowing down of the underlying oscillation in detection performance.

**Figure 6.**
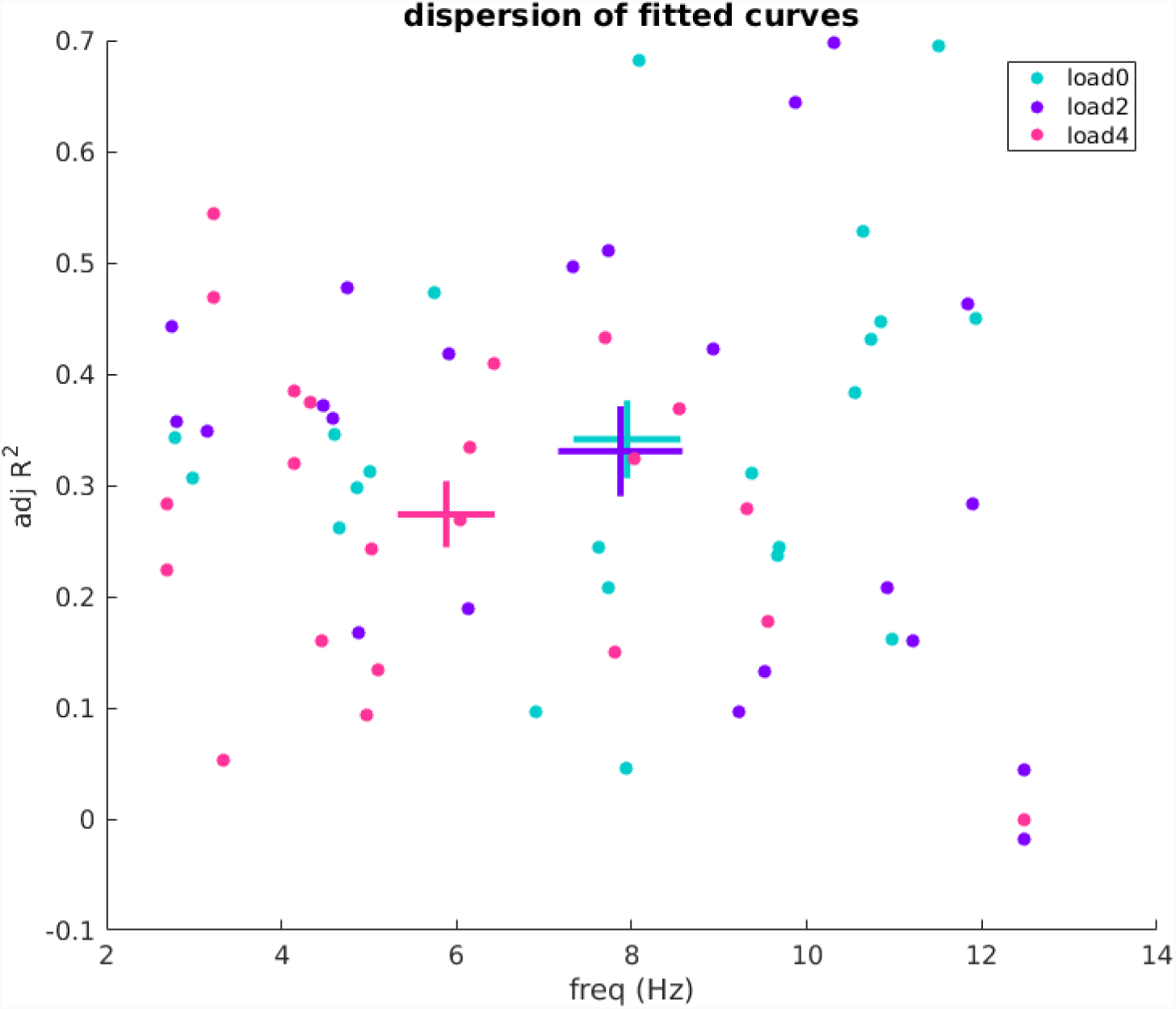
Average of HR time series after detrend with related sinusoidal fitting and significance testing: left insets show average trends, shaded areas show SEM, colors are coded as previously, (turquoise load0, violet load2, magenta load4), and the red line show the sinusoidal function fitted on data. The right insets show histograms of *adj R*^*2*^ obtained by the permutations in each specific condition, the black dashed line show the 95% percentile of the permuted distribution as threshold for significance, whereas the colored lines show where the observed *adj R*^*2*^ stands within the distribution of permuted values.

### 3.5 Spectral analysis of “good” vs “poor” performers

We performed the same spectral analysis described before but this time dividing the sample into “poor” and “good” performance based on Cowan’s K, and results are depicted in Figure 8. Load0 and load2 conditions, albeit showing local peaks, did not show any significant periodicities surpassing the 95° percentile the of permutation distribution.

**Figure 8:**
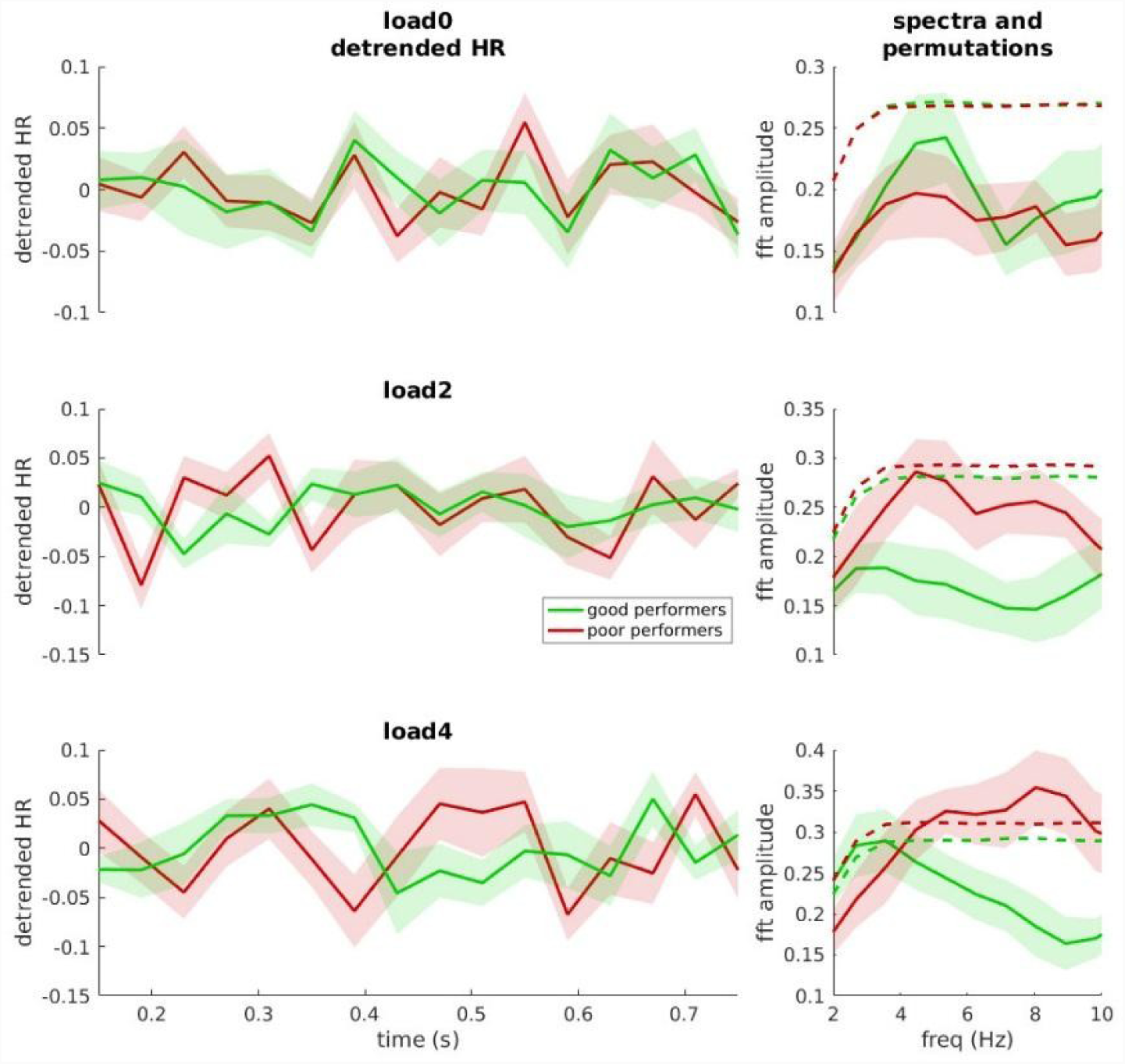
HR time series after detrend for “good” and “poor” performers, and average of correspective spectra: shaded areas show SEM, colors are coded as in Figure 3 and 4, (green good performers, red poor performers), dashed lines in the spectra insets show the 95th percentile of permutations.

In contrast, for the load4 condition a different scenario was found: the two samples of “good” and “poor” performers showed different spectra, where “good” performers show a maximum at ∼3.33 Hz (p_(uncorrected)_=.049) and the those with lower Cowan’s *k* showed a maximum at ∼7.5 Hz (p_(uncorrected)_=.004). It should be noted that when applying a more severe threshold for multiple comparison (see Methods), the significance for the peak at ∼3.33 Hz slightly exceeds the required alpha error probability (p_(corrected)_= .059), whereas the significance at ∼7.5 Hz for “poor” performers remains virtually unchanged (p_(corrected)_=.004). In any case, in the highest load condition, the two subgroups showed considerably different maxima and spectral signature, suggesting that the amount of items currently maintained in VWM (exemplified by Cowan’s K) was related to different rhythms in detection of a near threshold stimulus during the maintenance phase of VWM.

## 4. Discussion

In the present study we examined how loading VWM with different amounts of item (0, 2 or 4) to be maintained would impact on the detection of a near threshold target (flash) during the VWM maintenance phase. Consistent with previous research **(Kostantinou et al. 2012; Kostantinou et al. 2013)**, we found that increasing the load of VWM impaired detection of the flash during the memory maintenance phase. This first result suggests that the maintenance of items in VWM competes with the detection of stimuli in the external environment, a competition possibly linked to the active role of early visual areas in VWM information maintenance **(Harrison & Tong, 2009; Van Kerkoerle, Self & Roelfsema, 2017)**.

Our experiment further explored the trends of HR during the period of VWM maintenance by presenting the flash at several densely-sampled time points after the reset stimulus. This approach allowed us to uncover an oscillatory pattern at ∼5 Hz in flash detection when participants were given 4 items to maintain in VWM. This oscillatory pattern was shown by performing a spectral analysis of HR over time at the single subject level and also confirmed by the sinusoidal fitting over the group average of the same behavioural response. This result is in line with previous evidence showing that attention does not operate in a static fashion, but instead has an intrinsically oscillatory nature **(Busch & vanRullen, 2010; Fiebelkorn et al. 2011; Landau & Fries, 2012; Fiebelkorn et al. 2013; Fiebelkorn et al. 2018)**. Interestingly, when examining the goodness of sine fit at the level of single participants, we found that the best fitting frequency significantly shifted from the high theta-low alpha range (∼8 Hz) in low VWM load conditions (i.e. load0 and load2) to the theta range (∼5 Hz) in the high VWM load condition (i.e. load4). This result further supports our hypothesis that maintaining 4 items in VWM was related to a slowing down of the oscillatory patterns in detection, suggesting that the competition in early stages of visual processing between incoming and maintained stimuli can be resolved through an oscillation at ∼5 Hz. This rhythmic sampling in the theta range is in line with a growing number of studies showing that the division of attention among different spatial positions is associated with a division of a unitary sampling rhythm **(VanRullen et al., 2007; Landau & Fries, 2012; Fiebelkorn et al. 2013; VanRullen, 2018)**,

When splitting our sample of participants into “good” and “poor” performers based on their VWM performance, we observed two additional results that allow us to better characterize the influence that strategical attentional deployment to one task or the other might have on such oscillations in behavioural performance. First, we found a difference in the main frequency peaks for the flash detection between “good” and “poor” performers. Specifically, “good” performers had a slower (∼3.33 Hz) sampling frequency compared to “poor” performers (∼7.5 Hz) in the most burdening VWM condition (load4). Second, we found a drop in HR for “good” performers as compared to “poor” performers in load4. These results suggest that some participants may have strategically deployed more resources on the detection task, leading to higher sampling frequency, higher detection performance in terms of HR but lower VWM memory performance perhaps due to more shallow/quick periods of VWM maintenance. On the contrary, the “good” performers for the VWM task may have adapted a more “divided” strategy, leading on average to a higher amount of item stored in each trial, but at the same time to a slower sampling frequency for the detection task and, concurrently, to a poorer performance in the detection itself. This pattern provides support to the thesis that VWM and attention share common resources **(Kyonaga & Egner, 2013)**, possibly due to the centrality of an attentional sampling mechanism which strategically allocates resources for the task in use and the current goal. When the task is to pay attention to two -or more-spatial positions, the central attentional rhythm would be split **(VanRullen et al. 2007; Landau & Fries, 2012; Fiebelkorn et al. 2013)**, and the same seems to happen when these common resources are divided between two tasks, an internal (VWM) and an external (visual detection) task, as in the present study.

It should also be noted that the shared resource could have entailed the use of visual areas for the VWM task **(Harrison & Tong, 2009; Van Kerkoerle, Self & Roelfsema, 2017)**, with the behavioral oscillation reflecting a sort of multiplexing of those circuits for either top-down use (VWM maintenance) or bottom-up use (detection of the near threshold flash). Alternatively, the increase in theta frequency might have reflected a strategy to allocate resources to the VWM task, involving a slower oscillatory frequency, while prioritization of detection might have increased the influence of the faster 7-8 Hz frequency typically found in studies of visual selective attention.

Although the present study was behavioral, recent neuroimaging evidence may help to advance a plausible hypothesis about the neural mechanisms that might contribute to generate the oscillatory patterns in behaviour here outlined. For instance, converging evidence suggests that VWM maintenance relies on an interplay between frontal, parietal, and sensory cortices **(Christophel, Klink, Spitzer, Roelfsema, & Haynes, 2017)**, and that higher order areas influence the activity of sensory cortices by inter-areal coherence in the delta-theta band both in human **(Sauseng et al. 2009, Siebenhühner, Wang, Palva & Palva, 2016; Johnson, Dewar, Solbakk, Endestad, Meling, & Knight, 2017)** and in non-human primates **(Siegel, Warden, & Miller, 2009; Liebe, Hoerzer, Logothetis & Rainer, 2012)**. In this framework, this attentional sampling rhythm might act as a gatekeeper, implemented by higher order areas in order to parse the information incoming from sensory areas: when a competitive mechanism would be established, e.g. between two tasks relying on the same neural resources like in this case, the attentional sampling rhythm would slow down to allow both sources of information to alternatively access a particular computation, resolving in this way the competition. A comparable mechanism has been recently observed by Helfrich and colleagues **(Helfrich, Huang, Wilson & Knight, 2017)**, who showed that posterior alpha activity, entrained by either a predictive or an unpredicted sequence of stimuli, was modulated by a delta rhythm originating in frontal regions in the range of 2-4 Hz, selectively in the predictive condition. Such evidence suggests a role of higher order areas in top-down control of posterior alpha activity finalized to a facilitation of visual perception.

One limit of considering rhythmic sampling mechanism as a gatekeeper between incoming and maintained information is that the current study only shows evidence of sampling of external stimuli, i.e. we probed performance in visual detection over time of an external target, but we did not probe the actual maintenance of items internally. However, a recent study also using dense sampling of behavioural performance suggests that the temporal sampling mechanisms previously associated to attention occur even between objects maintained in VWM at ∼6 Hz **(Peters, Rahm, Kaiser, & Bledowski, 2018)**, closely resembling the periodicity at ∼ 5 Hz shown in the present paper.

Future work is needed to clarify the nature of a fluctuation between internal and external allocation of resources. One key question involves assessing whether the internal and external sampling of visual stimuli happens in antiphase, as might be expected when two tasks/locations alternate in terms of prioritization. Second, it would be useful to elucidate the relationship between behavioural sampling rhythm and the activity of fronto-parietal cortices in gating incoming information between incoming stimuli and maintained stimuli by the use of electrophysiological measures.

## 5. Conclusion

In summary, our results suggest that internally-driven VWM and externally-driven visual processing rely at least in part on shared attentional resources, since we found strong evidence for competition between the two tasks as exemplified by the reduction in performance in the detection of near threshold visual stimuli with the increase of VWM load. The overall pattern of results found here suggests that competition was resolved by a strategic allocation of attentional resources, which in our task was associated with an average ∼5 Hz sampling rhythm of the incoming visual stimulation in the full VWM load condition. Different strategies of attention allocation at the individual level, more focused either on the VWM task or on the detection task were linked to slower (∼3 Hz) or faster (∼7.5 Hz) sampling rhythms and to higher or lower VWM performance, respectively. All of these results converge in supporting the idea of a central sampling rhythm whose function is strategically pacing the flow of incoming visual information according to the current burden of the visual system due to the maintenance of visual information in memory.

## Acknowledgments

This research was supported by a European Research Council grant, “Construction of Perceptual Space-Time” (StG Agreement 313658).

